# Smile-to-Bert: a pre-trained Transformer model for molecular property prediction using SMILES representations

**DOI:** 10.1101/2024.10.31.621293

**Authors:** Maria Barranco-Altirriba, Vivian Würf, Enrico Manzini, Josch K. Pauling, Alexandre Perera-Lluna

**Affiliations:** B2SLab, Institute for Research and Innovation in Health (IRIS), Universitat Politècnica de Catalunya – BarcelonaTech, 08028 Barcelona, Spain; Networking Biomedical Research Centre in the subject area of Bioengineering, Biomaterials and Nanomedicine (CIBER-BBN), 28029 Madrid, Spain; Institut de Recerca Sant Joan de Déu (IRSJD), Esplugues de Llobregat, Barcelona, Spain; LipiTUM, Chair of Experimental Bioinformatics, TUM School of Life Sciences, Technical University of Munich, Maximus-von-Imhof-Forum 3, 85354 Freising, Germany; CioBio, Institute for Clinical Chemistry and Laboratory Medicine, University Hospital and Faculty of Medicine Carl Gustav Carus, Dresden University of Technology, 01307 Dresden, Germany

**Keywords:** BERT, SMILES, Physicochemical properties, binding affinity prediction

## Abstract

Molecular property prediction is crucial for drug discovery. Over the years, deep learning models have been widely used for these tasks; however, large datasets are often needed to achieve strong performances. Pre-training models on vast unlabeled data has emerged as a method to extract contextualized embeddings that boost performance on smaller datasets. The Simplified Molecular Input Line Entry System (SMILES) encodes molecular structures as strings, making them suitable for natural language processing. Transformers, known for capturing long-range dependencies, are well suited for processing SMILES. One such transformer-based architecture is Bidirectional Encoder Representations from Transformers (BERT), which only uses the encoder part of the Transformer and performs classification and regression tasks. Pre-trained transformer-based architectures using SMILES have significantly improved predictions on smaller datasets. Public data repositories such as PubChem, which provide SMILES, among other data, are essential for pre-training these models. SMILES embeddings that combine chemical structure and physicochemical property information could further improve performance on tasks such as Absorption, Distribution, Metabolism, Excretion, and Toxicity prediction. To this end, we introduce Smile-to-Bert, a pre-trained BERT architecture designed to predict 113 RDKit-computed molecular descriptors from PubChem SMILES. This model generates embeddings that integrate both molecular structure and physicochemical properties. We evaluate Smile-to-Bert on 22 datasets from the Therapeutics Data Commons and compare its performance with that of the 2-encoder model and a Transformer model. Smile-to-Bert achieves the best result on one dataset, while the combination of Smile-to-Bert with the other models leads to improved performance on 8 datasets. Additionally, the state-of-the-art Transformer is applied to Absorption, Distribution, Metabolism, Excretion, and Toxicity prediction for the first time, achieving the best performance on the Therapeutics Data Commons leaderboard of one dataset. Scientific Contribution: We present Smile-to-Bert, a model pre-trained to predict 113 molecular descriptors directly from SMILES representations. We evaluate our model on 22 property prediction datasets and compare the performance with other language models. We demonstrate that integrating our model with two other pre-trained models improves the performance in 8 datasets.

## 1 Introduction

Accurate prediction of molecular properties [1] plays a crucial role in computer-aided drug discovery. In recent years, there has been a significant increase in the use of deep learning models for drug discovery, driven by the increasing availability of compound-related data [2]. Public databases such as PubChem [3], which contain large amounts of chemical information, exemplify the increasing availability of data in the field. However, acquiring large labeled datasets for molecular property prediction is both challenging and costly, limiting the training of deep learning models [1, 4]. Transfer learning has emerged as a solution to overcome data limitations in deep learning [4, 5, 6]. It involves pre-training a model on one data domain and then adapting it to a different domain of interest, which allows the extraction of meaningful features from vast datasets, even when relevant labels are limited [4].

The Simplified Molecular Input Line Entry System (SMILES) are text-based representations that encode a molecular graph in a simple, human-readable sequence of characters [7, 8]. Due to their linear nature, SMILES are widely used in natural language processing (NLP) algorithms [9]. The order of SMILES characters does not always represent the physical proximity of atoms in a molecule, which can complicate the understanding of long-range dependencies. This poses a challenge for recurrent neural networks and showcases the usefulness of transformer-based architectures [4].

The Transformer, first introduced by Vaswani et al. [10], uses a self-attention mechanism that processes all inputs simultaneously, thus learning long-range dependencies as easily as short-range ones. Using self-attention to process SMILES strings allows for the learning of atomic structure features without relying on assumptions about the sequential nature of the data. Transfer learning has been widely used in Transformer-based architectures. Some decoder-only architectures use unidirectional language models during pre-training, which may limit their ability to learn effectively for certain tasks, such as sentence-level understanding, where context from both directions is crucial. To address these limitations, Bidirectional Encoder Representations from Transformers (BERT), which only uses the encoder part of the Transformer, was introduced. In the original paper, BERT enables bidirectional context understanding using a masked language modeling as a pre-training objective [11].

BERT architectures have been pre-trained with the goal of generating contextualized embeddings that can boost the performance of models trained on smaller datasets [5]. More specifically, transformer-based architectures pre-trained through various tasks have been applied to a wide range of applications for processing SMILES. For instance, to address the limited amount of data in binding affinity assays, a Transformer architecture was pre-trained on the task of translating SMILES strings to International Union of Pure and Applied Chemistry (IUPAC) names. By leveraging the SMILES embeddings obtained by the pre-trained model, binding affinity prediction showed improvement across three datasets compared to using untrained embedding vectors [4]. Similarly, transformer-based architectures have been pre-trained using a masked language modeling (MLM) objective with SMILES and then fine-tuned to predict molecular properties [1, 12, 13]. Zheng and Tomiura [14] proposed a 2-encoder BERT that improved token recovery in an MLM task. The first encoder takes the complete SMILES string as input, while the second encoder processes the masked SMILES along with the classification token embedding from the first encoder. This model was then fine-tuned to predict molecular properties across 22 datasets from Therapeutics Data Commons (TDC) [15, 16], achieving better performance than a model pre-trained solely on the MLM task. TDC provides AI-ready datasets and tasks for therapeutic machine learning, along with a set of tools for easy benchmarking of algorithms. Among the TDC datasets, 22 Absorption, Distribution, Metabolism, Excretion and Toxicity (ADMET) datasets are available for predicting and classifying small molecule properties [16].

Understanding molecules’ behavior in humans or the environment largely depends on their key physicochemical properties. These properties can currently be measured through experimental methods or predicted using quantitative structure-property relationship (QSPR) models. However, experimental methods are often costly, time-consuming, and challenging. Consequently, in silico QSPR modeling provides a valuable alternative by enabling the rapid estimation of physicochemical properties through the identification of relationships between molecular structures and their associated properties [17]. Furthermore, a random forest trained with fingerprints and 200 molecular descriptors calculated with RDKit shows very good performance in ADMET datasets[18], showing the importance of molecular descriptors in these tasks.

Given this context, we hypothesize that incorporating physicochemical properties information into embedding vectors might enhance their richness and utility for various SMILES-related tasks, such as ADMET properties prediction. Therefore, we present Smile-to-Bert, a BERT architecture pre-trained on a large subset of Pub-Chem molecules to predict 113 molecular descriptors computed with RDKit from SMILES. In this work, we (1) pre-train a BERT architecture by predicting 113 molecular descriptors from SMILES (2) interpret the pre-trained model and its predictions using integrated gradients, (3) fine-tune our pre-trained model to predict ADMET properties with 22 TDC datasets, (4) compare our results with the Transformer from Morris et al. [4] and the 2-encoder model from Zheng and Tomiura [14] and (5) combine both models with Smile-to-Bert to improve the overall performance.

## 2 Methods

### 2.1 Pre-training data and model architecture

A set of sdf PubChem files available in https://ftp.ncbi.nlm.nih.gov/pubchem/ Compound/CURRENT-Full/SDF/ were downloaded using the RCurl R package v1.98.1.14 [19] and the SMILES were extracted with ChemmineR package v3.46.0 [20]. RDKit v2024.09.4 was used to compute the 200 molecular descriptors used in MapLight [18]. Compounds were discarded based on several filters. Figure 1 shows the general scheme of Smile-to-Bert pre-training.

**Fig. 1.**
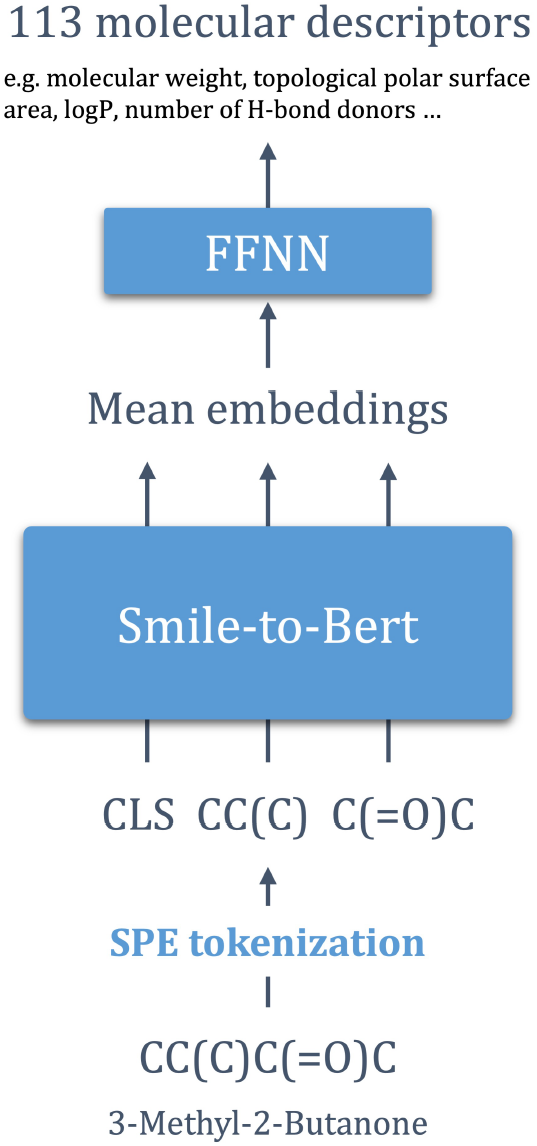
Smile-to-Bert pre-training model. Smile-to-Bert pre-training model. SPE stands for SMILES Pair Encoding, and FFNN denotes a feed-forward neural network.

Specific details of the input, model architecture and output are described in the following subsections.

#### 2.1.1 Input - SMILES

The first step in NLP algorithms is tokenizing the sequence. Various approaches exist for tokenizing SMILES, but character-level tokenization can lead to issues, such as splitting atoms represented by multiple characters into separate tokens. Atom-level tokenization is a more common approach that ensures each atom is extracted as a distinct token, regardless of the number of characters used to represent that atom. A more sophisticated method is SMILES pair encoding (SPE), a substructural tokenization algorithm inspired by the byte pair encoding algorithm. SPE detects and preserves common SMILES substrings as unique tokens, generating tokens with richer information that more accurately reflect molecular functionalities compared to atom-level tokenization. Additionally, input sequences become shorter compared to those produced by atom-level tokenization [8]. Therefore, we employed the SPE_tokenizer using SPE Tokenizer from SmilesPE module [8].

All SMILES sequences were limited to a maximum length of 100 tokens.

#### 2.1.2 Model Architecture

In a BERT model, the encoder section of the Transformer is employed to generate contextualized representations for each input token, which can then be used for downstream tasks, such as predicting a vector of physicochemical properties. Initially, each token in the input sequence is mapped to a dense embedding vector, capturing its semantic meaning. A positional encoding is added to these embeddings to incorporate the sequential information of the tokens. These token embeddings are then passed through a series of stacked encoder layers. Each encoder layer applies multi-head selfattention, which allows capturing complex dependencies between tokens. After this, feed-forward neural networks (FFNN) and normalization operations are applied. The final contextualized vector serves as input for the specific downstream task.

Our BERT architecture was composed of 4 encoder layers, 8 attention heads, and an embedding size of 512 was used. We used a reduced version of the original BERT to speed up the training. A scheme of the architecture can be seen in Figure S1. A sequence length of 100 was enforced by padding the token sequences. The contextualized embeddings that correspond to non-padding tokens were averaged, and the final embedding vector of size 512 was used to predict 113 molecular descriptors, using one linear layer.

#### 2.1.3 Output - Molecular descriptors

The 200 molecular descriptors used in MapLight [18] were computed using RDKit for 3,976,178 SMILES. Molecular descriptors with identical first and third quartiles were removed, reducing the feature set by 86. Additionally, the Ipc descriptor, which quantifies the information content of a molecule’s structure, was discarded because its values fell outside the range [-1e6, 1e6] for more than 90% of the molecules. After filtering out these descriptors, molecules with any remaining descriptor values outside this range were also removed, yielding a final dataset of 3,976,176 molecules. Before training, properties values were normalized by subtracting their corresponding median value and dividing by their IQR. This scaling was selected because of its robustness regardless the presence of outliers. Once the properties were scaled, they were multiplied by a constant factor of 100. This last step was performed to avoid the vanishing gradients problem (see Supporting Information for details).

### 2.2 Model training

The models were trained using PyTorch v2.3.0+cu121 [21], and experiments were tracked with wandb v0.16.3 [22]. L1Loss function was used to compute the loss. The Adam optimizer [23] was employed with a weight decay of 0.01. The initial learning rate was set to 2e-4 for the BERT encoder and 5e-5 for the final FFNN responsible for predicting molecular descriptors, thereby encouraging the BERT model to train more effectively. The optimizer scheduler used by Vaswani et al [10] was employed with 2000 warm-up steps. Data was split randomly for training (95%) and validation (5%) and a batch size of 64 was used during training. The models were trained for 20 epochs and the models which obtained the minimum total loss in validation were selected. Accelerate v0.27.2 [24] was used for distributed training in a total number of 2 virtual machines with 2 NVIDIA A40 GPUs each.

### 2.3 Model performance and interpretation

Learning curves for each molecular descriptor were created. Additionally, a Principal Component Analysis (PCA) of the embeddings of 79,524 randomly selected SMILES was conducted using sklearn v1.4.0 [25]. The first two principal components were used to visualize the selected embeddings colored by six molecular descriptors.

### 2.4 ADMET datasets prediction

The usefulness of the models was evaluated using the 22 ADMET datasets available in TDC. Each model consisted of a pre-trained Encoder and a FFNN, which took the mean of the non-padding tokens from the contextualized embedding matrix as input. The FFNN, used in the work of Zheng and Tomiura [14], contains two layers: one with dimensions *embedding size* × *hidden layer* and the other with dimensions *hidden layer* × 1. It applies a ReLU activation function to the output of the first layer and a sigmoid function in the final output in classification cases. Also, a dropout of 0.1 is applied in all layers. For Smile-to-Bert and the Transformer from Morris et al. [4], *embedding size* was 512, and *hidden layer* was tested with values of 256 and 512. For the 2-encoder model [14], *embedding size* was 256, and *hidden layer* was 256. In all cases, the models were tested with two strategies: freezing all layers except the last one and freezing all layers except the last two. In addition to these models, the combination of models was also evaluated to assess the usefulness of integrating molecular descriptor information into embeddings obtained from unsupervised tasks. The combination was performed by concatenating the contextualized embeddings of each model and inputting the resulting matrix into the FFNN. The 2-encoder model was combined with Smile-to-Bert, resulting in an FFNN with an *embedding size* of 768, and two *hidden layer* values (256 and 768) were tested. The Transformer was also combined with Smile-to-Bert, resulting in an FFNN with an *embedding size* of 1024, and two *hidden layer* values (256 and 512) were tested.

The Python package pyTDC provides a set of tools for evaluating models under identical conditions, including scaffold splits for training, validation, and test sets, as well as validation metrics for regression (Mean Absolute Error (MAE) and Spearman correlation) and classification (Area Under the Receiver Operating Characteristic Curve (AUROC) and Area Under the Precision-Recall Curve (AUPRC)) specific to each dataset. The validation strategy involved splitting the training-validation data into training and validation sets using the five seeds indicated by TDC. The model was trained with early stopping, allowing a maximum of 10 epochs for validation loss to decrease. The best-performing model, selected based on validation loss, was then used for testing. Data preparation for each dataset involved SMILES tokenization, using each model-specific approach, and output scaling via z-score computation for regression datasets. The mean and standard deviation of the output were computed using only the training data.

BCELoss and MSELoss functions were used for classification and regression tasks, respectively. The Adam optimizer [23] was employed with a learning rate of 1e-4. The models were trained for a maximum number of 50 epochs.

## 3 Results

### 3.1 Data description

The descriptive statistics of a random selection of 15 RDKit-computed molecular descriptors are shown in Table 1, providing an overview of the data used for pretraining.

**Table 1.** Descriptive statistics of 15 randomly selected molecular descriptors. A total number of 3,976,176 compounds were used for pre-training.

|  | Mean | SD | Min | 25% | 50% | 75% | Max |
| --- | --- | --- | --- | --- | --- | --- | --- |
| SlogP_VSA10 | 5.2563 | 7.0639 | 0.0000 | 0.0000 | 4.7945 | 5.9483 | 447.8223 |
| Chi0v | 13.3166 | 4.5910 | 0.4082 | 10.7247 | 13.0526 | 15.1330 | 198.7401 |
| MolMR | 101.5520 | 35.1565 | 0.0000 | 81.6435 | 99.1497 | 116.2480 | 1510.2524 |
| RingCount | 2.9990 | 1.4497 | 0.0000 | 2.0000 | 3.0000 | 4.0000 | 63.0000 |
| Chi3n | 2.8461 | 1.2248 | 0.0000 | 2.1082 | 2.7344 | 3.4080 | 52.1561 |
| NumAliphaticRings | 0.8888 | 1.1132 | 0.0000 | 0.0000 | 1.0000 | 1.0000 | 63.0000 |
| SMR_VSA1 | 45.7416 | 21.7184 | 0.0000 | 33.2823 | 43.0017 | 53.6643 | 1312.5082 |
| PEOE_VSA7 | 39.7303 | 20.5908 | 0.0000 | 24.9791 | 37.0640 | 49.8195 | 829.6910 |
| NumAromaticHeterocycles | 0.6689 | 0.8207 | 0.0000 | 0.0000 | 0.0000 | 1.0000 | 18.0000 |
| Chi3v | 3.3808 | 1.8915 | 0.0000 | 2.4260 | 3.2029 | 4.0862 | 655.5112 |
| VSA_EState7 | -11.3178 | 18.9909 | -1379.8901 | -14.2587 | -5.6605 | -0.9398 | 12.5000 |
| MinAbsPartialCharge | 0.2788 | 0.0749 | 0.0000 | 0.2369 | 0.2771 | 0.3323 | 6.0000 |
| SlogP_VSA2 | 33.7954 | 23.6502 | -0.8818 | 19.0683 | 29.8187 | 42.9626 | 2170.1561 |
| fr_halogen | 0.6583 | 1.3189 | 0.0000 | 0.0000 | 0.0000 | 1.0000 | 102.0000 |
| EState_VSA9 | 34.5672 | 14.0227 | 0.0000 | 25.9054 | 32.9586 | 41.2436 | 911.7682 |

### 3.2 Model performance and interpretation

The learning curves for each molecular descriptor are shown in the Results section of the Supporting Information.

Regarding the interpretation of the model, Figure 2 shows the first two principal components of 79,524 randomly selected SMILES embeddings generated using Smile-to-Bert. The plots are color-coded by six molecular descriptors: ExactMolWt, representing the exact molecular weight; MinAbsEStateIndex, the minimum absolute electrotopological state index; qed, the quantitative estimate of drug-likeness; FpDensityMorgan1, the density of Morgan fingerprint features at radius 1; MaxEStateIndex, the maximum electrotopological state index; and FractionCSP3, the fraction of sp^3^-hybridized carbon atoms.

**Fig. 2.**
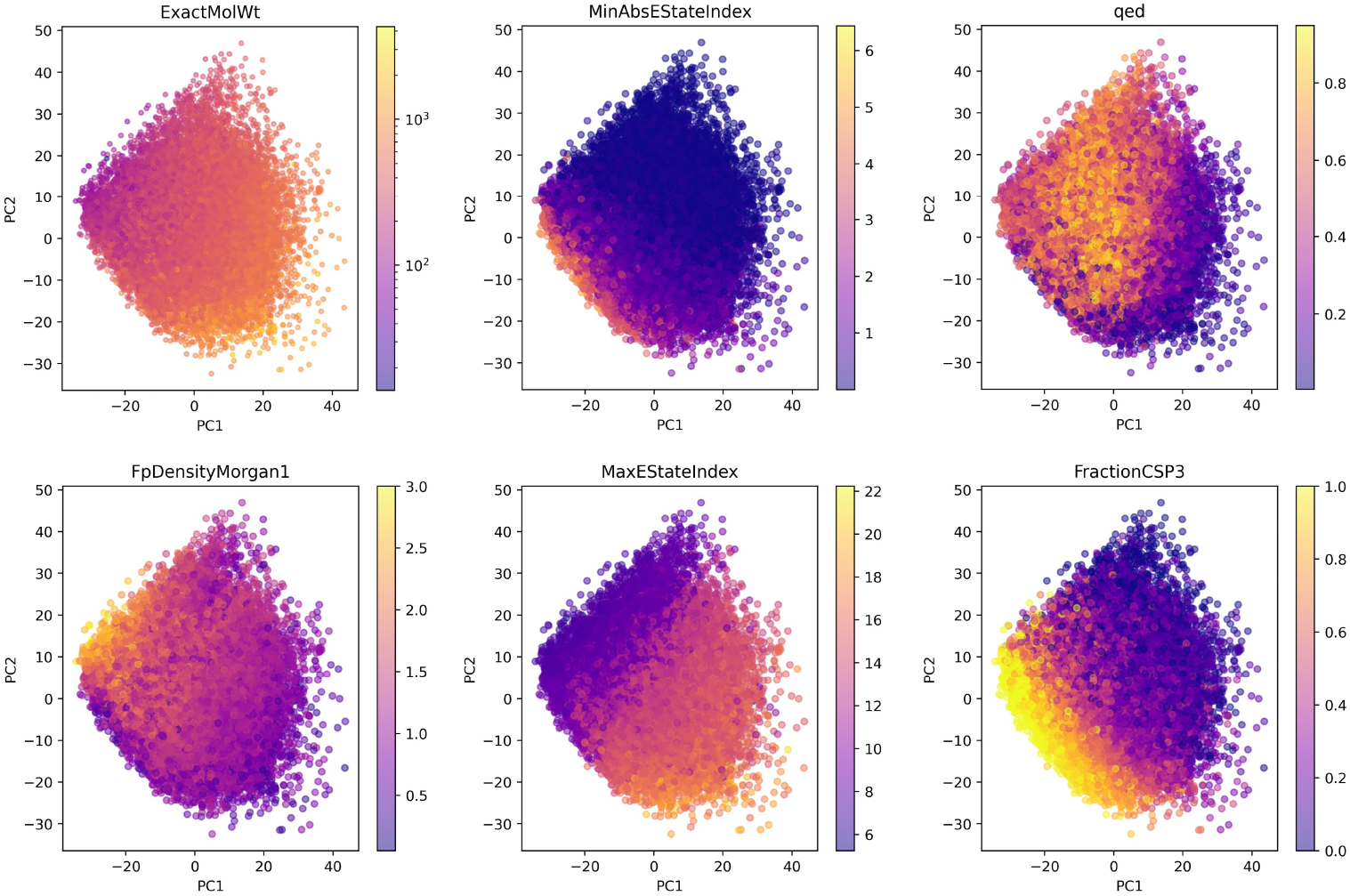
PCAs of the embeddings colored by six molecular descriptors.

### 3.3 ADMET datasets prediction

Table 2 shows the results for Smile-to-Bert, the Transformer model from Morris et al. [4], and the 2-encoder model [14] as individual models, while Table 3 presents the results for combinations of the 2-encoder model and the Transformer with Smile-to-Bert. Of the 22 datasets, the Transformer achieves the best performance in 9, the 2-encoder model in 5, and Smile-to-Bert in 1. The combined 2-encoder and Smile-to-Bert model achieves the best performance in 7 datasets, while the Transformer and Smile-to-Bert combination performs best in 1 dataset.

**Table 2.** Evaluation results for individual models using TDC datasets. Values represent the mean of the metric (standard deviation). The best results across all configurations are highlighted in bold.

| Datasets | ADMET | Size | Metric | Smile-to-Bert | 2-encoder 1L | 2-encoder 2L | Transf. 1L256 | Transf. 1L512 | Transf. 2L512 |
| --- | --- | --- | --- | --- | --- | --- | --- | --- | --- |
| caco2_wang | Absorption | 910 | MAE | 0.347 (0.011) | 0.382 (0.020) | 0.423 (0.019) | 0.326 (0.013) | 0.326 (0.017) | **0.314 (0.008)** |
| hia_hou | Absorption | 578 | ROC-AUC | 0.966 (0.006) | 0.984 (0.004) | 0.973 (0.011) | 0.988 (0.003) | **0.991 (0.002)** | 0.990 (0.002) |
| pgp_broccatelli | Absorption | 1218 | ROC-AUC | 0.837 (0.018) | **0.932 (0.005)** | **0.932 (0.004)** | 0.911 (0.013) | 0.915 (0.008) | 0.921 (0.013) |
| bioavailability_ma | Absorption | 640 | ROC-AUC | 0.622 (0.022) | 0.635 (0.041) | 0.603 (0.047) | 0.673 (0.021) | **0.677 (0.020)** | 0.673 (0.022) |
| lipophilicity_astrazeneca | Absorption | 4200 | MAE | 0.869 (0.011) | 0.621 (0.006) | 0.628 (0.010) | **0.579 (0.006)** | 0.591 (0.019) | 0.582 (0.009) |
| solubility_aqsoldb | Absorption | 9982 | MAE | 1.187 (0.082) | 0.937 (0.018) | 0.929 (0.029) | 0.875 (0.044) | 0.857 (0.047) | **0.835 (0.016)** |
| bbb_martins | Distribution | 2030 | ROC-AUC | 0.855 (0.022) | 0.862 (0.034) | 0.863 (0.038) | 0.898 (0.011) | **0.905 (0.009)** | 0.898 (0.008) |
| ppbr_az | Distribution | 2790 | MAE | 11.307 (0.496) | 9.796 (0.330) | 9.578 (0.286) | 9.066 (0.776) | 9.038 (0.760) | **8.769 (0.383)** |
| vdss_lombardo | Distribution | 1130 | Spearman | 0.402 (0.107) | 0.233 (0.088) | 0.232 (0.032) | 0.458 (0.035) | **0.471 (0.038)** | 0.430 (0.009) |
| cyp2d6_veith | Metabolism | 13130 | PR-AUC | 0.502 (0.011) | 0.652 (0.006) | 0.657 (0.005) | 0.639 (0.004) | 0.631 (0.007) | 0.645 (0.012) |
| cyp3a4_veith | Metabolism | 12328 | PR-AUC | 0.733 (0.016) | 0.855 (0.004) | **0.858 (0.003)** | 0.831 (0.007) | 0.834 (0.006) | 0.831 (0.003) |
| cyp2c9_veith | Metabolism | 12092 | PR-AUC | 0.641 (0.018) | 0.765 (0.006) | 0.765 (0.005) | 0.744 (0.008) | 0.745 (0.011) | 0.747 (0.005) |
| cyp2d6_substrate_carbonmangels | Metabolism | 667 | PR-AUC | 0.559 (0.013) | **0.689 (0.042)** | 0.681 (0.032) | 0.645 (0.013) | 0.645 (0.017) | 0.651 (0.008) |
| cyp3a4_substrate_carbonmangels | Metabolism | 670 | ROC-AUC | 0.532 (0.025) | **0.646 (0.028)** | 0.617 (0.040) | 0.594 (0.024) | 0.594 (0.021) | 0.587 (0.007) |
| cyp2c9_substrate_carbonmangels | Metabolism | 669 | PR-AUC | 0.361 (0.018) | 0.386 (0.036) | 0.392 (0.039) | 0.365 (0.032) | 0.372 (0.021) | 0.378 (0.016) |
| half_life_obach | Excretion | 667 | Spearman | **0.363 (0.065)** | 0.033 (0.063) | 0.045 (0.050) | 0.304 (0.032) | 0.242 (0.031) | 0.259 (0.053) |
| clearance_microsome_az | Excretion | 1102 | Spearman | 0.432 (0.027) | 0.585 (0.031) | 0.600 (0.034) | 0.543 (0.010) | 0.566 (0.025) | 0.561 (0.026) |
| clearance_hepatocyte_az | Excretion | 1213 | Spearman | 0.318 (0.019) | 0.399 (0.049) | 0.426 (0.049) | 0.418 (0.017) | 0.404 (0.029) | 0.405 (0.040) |
| herg | Toxicity | 655 | ROC-AUC | 0.753 (0.016) | 0.830 (0.009) | 0.826 (0.010) | 0.838 (0.005) | 0.847 (0.004) | **0.852 (0.013)** |
| ames | Toxicity | 7278 | ROC-AUC | 0.775 (0.019) | 0.832 (0.005) | 0.826 (0.006) | 0.817 (0.007) | 0.815 (0.003) | 0.812 (0.008) |
| dili | Toxicity | 475 | ROC-AUC | 0.839 (0.019) | 0.791 (0.126) | 0.855 (0.027) | 0.865 (0.020) | 0.862 (0.020) | 0.860 (0.029) |
| ld50_zhu | Toxicity | 7385 | MAE | 0.678 (0.017) | 0.653 (0.019) | 0.654 (0.013) | 0.666 (0.023) | 0.655 (0.016) | 0.674 (0.013) |

**Table 3.** Evaluation results for combined models using TDC datasets. Values represent the mean of the metric (standard deviation). The best results across all configurations are highlighted in bold.

| Datasets | ADMET | Size | Metric | Combined 2-encoder 1L768 | Combined 2-encoder 2L768 | Combined Transf. 2L512 |
| --- | --- | --- | --- | --- | --- | --- |
| caco2_wang | Absorption | 910 | MAE | 0.334 (0.037) | 0.336 (0.029) | 0.361 (0.017) |
| hia_hou | Absorption | 578 | ROC-AUC | 0.971 (0.009) | 0.968 (0.012) | 0.973 (0.012) |
| pgp_broccatelli | Absorption | 1218 | ROC-AUC | 0.882 (0.042) | 0.906 (0.013) | 0.878 (0.027) |
| bioavailability_ma | Absorption | 640 | ROC-AUC | 0.620 (0.016) | 0.625 (0.047) | 0.632 (0.046) |
| lipophilicity_astrazeneca | Absorption | 4200 | MAE | 0.604 (0.023) | 0.604 (0.009) | 0.642 (0.018) |
| solubility_aqsoldb | Absorption | 9982 | MAE | 0.849 (0.015) | 0.838 (0.020) | 0.908 (0.042) |
| bbb_martins | Distribution | 2030 | ROC-AUC | 0.879 (0.024) | 0.885 (0.036) | 0.878 (0.026) |
| ppbr_az | Distribution | 2790 | MAE | 9.417 (0.218) | 9.660 (0.328) | 9.558 (0.423) |
| vdss_lombardo | Distribution | 1130 | Spearman | 0.184 (0.132) | 0.235 (0.078) | 0.343 (0.104) |
| cyp2d6_veith | Metabolism | 13130 | PR-AUC | 0.677 (0.012) | **0.679 (0.009)** | 0.631 (0.010) |
| cyp3a4_veith | Metabolism | 12328 | PR-AUC | 0.857 (0.001) | 0.854 (0.008) | 0.821 (0.010) |
| cyp2c9_veith | Metabolism | 12092 | PR-AUC | **0.768 (0.007)** | 0.767 (0.009) | 0.722 (0.009) |
| cyp2d6_substrate_carbonmangels | Metabolism | 667 | PR-AUC | 0.676 (0.029) | 0.684 (0.032) | 0.598 (0.013) |
| cyp3a4_substrate_carbonmangels | Metabolism | 670 | ROC-AUC | 0.616 (0.016) | 0.607 (0.029) | 0.522 (0.020) |
| cyp2c9_substrate_carbonmangels | Metabolism | 669 | PR-AUC | 0.397 (0.015) | 0.394 (0.018) | **0.400 (0.017)** |
| half_life_obach | Excretion | 667 | Spearman | 0.244 (0.056) | 0.231 (0.083) | 0.343 (0.088) |
| clearance_microsome_az | Excretion | 1102 | Spearman | 0.606 (0.034) | **0.621 (0.013)** | 0.531 (0.035) |
| clearance_hepatocyte_az | Excretion | 1213 | Spearman | **0.448 (0.024)** | 0.401 (0.041) | 0.398 (0.036) |
| herg | Toxicity | 655 | ROC-AUC | 0.843 (0.022) | 0.834 (0.029) | 0.819 (0.026) |
| ames | Toxicity | 7278 | ROC-AUC | 0.830 (0.009) | **0.833 (0.005)** | 0.803 (0.008) |
| dili | Toxicity | 475 | ROC-AUC | 0.876 (0.017) | **0.891 (0.017)** | 0.861 (0.019) |
| ld50_zhu | Toxicity | 7385 | MAE | **0.633 (0.007)** | **0.633 (0.020)** | 0.659 (0.014) |

Only the model configurations that achieved the best results in at least one dataset are included in the tables. For Smile-to-Bert, the configuration with the last two layers trained and a hidden layer size of 512 yielded the best result for one dataset. For all other models, the configuration is labeled as NLX, where N represents the number of trained layers and X denotes the hidden layer size.

Table 3 presents the results obtained from combining the 2-encoder and Transformer models with Smile-to-Bert.

## 4 Discussion

In this study, we have pre-trained a BERT architecture to predict 113 molecular descriptors from SMILES and utilized the pre-trained encoder, which contains molecular knowledge, to predict ADMET properties across 22 datasets. We refer to this model as Smile-to-Bert and have compared our results with those from two other language models. Additionally, we have combined these models with Smile-to-Bert, achieving better performance in some cases.

The Smile-to-Bert learning curves for each molecular descriptor in our dataset are shown in Figures S5-S11. Despite the presence of extreme outliers in some of the characteristics (see Table 1), Smile-to-Bert can predict 113 molecular descriptors from SMILES with high accuracy, achieving this in just a few epochs. This demonstrates that attention-based deep learning architectures have strong potential to efficiently understand the molecular structures of SMILES.

Figure 2 presents the first two principal components of randomly selected SMILES embeddings, colored according to six molecular descriptors. Two of these descriptors are measures of electrotopological state (EState) indices, which combine electronic and topological characteristics [26]. Previous studies have shown that atom-type EState indices contribute significantly to modeling aqueous solubility (logS), lipophilicity (logP), and toxicity [27]. The Quantitative Estimate of Drug-likeness (QED) was empirically derived from approved drug data, considering eight widely used molecular properties known to influence drug attrition, including molecular weight [28]. Another measure of drug-likeness is the fraction of sp^3^-hybridized carbon atoms, which has been linked to drug success rates [29]. Additionally, the density of Morgan fingerprints provides insight into molecular complexity, and fingerprint-based random forest models have demonstrated promising results in ADMET prediction [30]. A gradient of these properties can be observed in Figure 2. The fact that our embeddings capture information about these key molecular descriptors, which are directly relevant to drug development, suggests their potential utility in ADMET property prediction.

We evaluated a set of models using 22 datasets for ADMET property prediction. According to the results shown in Tables 2 and 3, the Transformer, as an individual model, performs better than the others. However, the 2-encoder model, whether used individually or in combination, appears more frequently in the top-performing outcomes. In contrast, Smile-to-Bert by itself only achieves the best result in one dataset, but it improves the performance of the other two models in 8 cases. This supports the hypothesis that integrating molecular descriptor knowledge into these prediction tasks can enhance overall accuracy, as previously suggested by Notwell and Wood [18]. However, despite improving the performance of other models, Smile-to-Bert still performs worse than both the Transformer and the 2-encoder model. We believe this is due to the pre-training tasks used. The Transformer is pre-trained to translate SMILES into IUPAC names, and the 2-encoder model is trained to predict masked tokens, likely resulting in embeddings that capture general chemical information. In contrast, the embeddings from Smile-to-Bert are specifically tailored to predict a set of physicochemical properties. Although the number of properties predicted is large, this training task likely results in very specific embedding vectors that may not generalize as well as broader ones. However, if the new dataset closely relates to some of the molecular descriptors from our dataset, performance on the new dataset could still be strong, as might be the case with the ‘half_life_obach’_dataset.

In addition, certain patterns can be observed regarding the ADMET property predicted and the best-performing model. The Transformer model performs best for absorption and distribution. For metabolism, models that include the 2-encoder architecture, either alone or in combination with Smile-to-Bert, achieve the highest performance. For excretion, Smile-to-Bert is involved in achieving the best performance in the three datasets, suggesting the importance of molecular descriptors in predicting this property. Finally, the combination of Smile-to-Bert and the 2-encoder model performs best for toxicity prediction. Further analysis of this trend is important, either to develop a model that combines the best features of all three and performs well across all datasets or to design specialized models for specific properties.

The Therapeutics Data Commons (TDC) [15, 16] provides the opportunity to compare results under identical conditions across a wide range of datasets, and researchers can present their results on TDC leaderboards. When comparing the results from this study with those publicly available on the leaderboards, the Transformer outperforms all other algorithms in the ‘hia_hou’ dataset, while the 2-encoder model ranks third in the ‘pgp_broccatelli’ dataset. This is promising, considering none of the models were specifically fine-tuned or designed for these tasks. However, for the remaining datasets, the results tend to fall in the middle or lower end of the table. As previously mentioned, a random forest using fingerprints and 200 molecular descriptors consistently ranks at the top. This could be attributed to the type of input used, the simplicity of the model, or the ease of fine-tuning its parameters, or a combination of these factors. Nevertheless, language models have demonstrated impressive capabilities across a wide range of NLP tasks, and there is no reason to believe they would not exhibit similar performance in predicting molecular properties. Therefore, further research is needed to fully leverage the potential of attention-based pre-trained models capable of learning optimal molecular features for accurate prediction of molecular properties. This study has several strengths, including the extensive data used for pre-training and validation, the low losses achieved during pre-training and the full availability of the code. Additionally, we evaluate 5 attention-based language models, 3 individuals and 2 combined, to predict 22 ADMET properties in TDC datasets from SMILES. From these models, the Transformer from Morris et al. [4] is used to predict these properties for the first time, showing promising results, and obtaining the best result in the TDC leaderboards for one dataset. Finally, we integrate both the Transformer and the 2-encoder model from Zheng and Tomiura [14] with Smile-to-Bert, achieving better results in 8 datasets. This showcases the model’s potential in integrative approaches. However, there are some limitations. First, Smile-to-Bert does not show stronger performances in most of the datasets, probably due to the supervised pretraining task used. Moreover, simpler models like random-forest outperform the results that we obtain in most of the datasets, highlighting the importance of further research in this topic.

## 5 Conclusion

In this work, we present Smile-to-Bert, a BERT architecture pre-trained to predict 113 molecular descriptors from SMILES, generating contextualized embeddings that capture molecular features known to be important for property prediction in drug discovery. We evaluate our model, along with two other pre-trained language models, on 22 ADMET property datasets from the TDC. Smile-to-Bert achieves the best performance on one dataset, while the other models deliver competitive results on two TDC leaderboards. Additionally, we combine Smile-to-Bert with the 2-encoder and Transformer models, achieving superior performance on 8 datasets, highlighting the potential of integrative approaches with language models. Our findings suggest that a model pre-trained with an unsupervised or semi-supervised task, that incorporates molecular descriptors knowledge could generate richer embeddings that enhance performance in other tasks. The code for pre-training and interpreting Smile-to-Bert can be accessed at https://github.com/m-baralt/smile-to-bert, and the code for predicting TDC datasets using the five models analyzed in this paper is available at https://github.com/m-baralt/smile-to-bert-tdc.

## Declarations

### 5.1 Availability of data and materials

The code and data used for this paper are available at https://github.com/m-baralt/ smile-to-bert and https://github.com/m-baralt/smile-to-bert-tdc.

### 5.4 Authors’ contributions

MBA, VW, EM, JKP and APL conceived and designed the study. MBA developed the algorithm, obtained the results, and drafted the manuscript. VW, EM, JKP, and APL provided expert review throughout the project. JKP and APL supervised the study and contributed funding and access to computational resources. All authors have reviewed and approved the final version of the manuscript.

#### 5.5 Acknowledgements

This work was supported by the de.NBI Cloud within the German Network for Bioinformatics Infrastructure (de.NBI) and ELIXIR-DE (Forschungszentrum Jülich and W-de.NBI-001, W-de.NBI-004, W-de.NBI-008, W-de.NBI-010, W-de.NBI-013, W-de.NBI-014, W-de.NBI-016, W-de.NBI-022). The first author gratefully acknowledges the Universitat Polit`ecnica de Catalunya and Banco Santander for the financial support of her predoctoral grant.

### 5.6 Consent to Publish

Not applicable.

### 5.7 Ethics and Consent to Participate

Not applicable.

#### 6 Abbreviations

SMILES: Simplified Molecular Input Line Entry System
BERT: Bidirectional Encoder Representations from Transformers
ADMET: Absorption, Distribution, Metabolism, Excretion, and Toxicity
TDC: Therapeutics Data Common
NLP: Natural Language Processing
IUPAC: International Union of Pure and Applied Chemistry
QSPR: Quantitative Structure-Property Relationship
SPE: SMILES pair encoding
IQR: Interquartile range
PCA: Principal Component Analysis
FFNN: Feed-Forward Neural Network
MAE: Mean Absolute Error
AUPRC: Area Under the Precision-Recall Curve
AUROC: Area Under the Receiver Operating Characteristic Curve

