## Supporting Information for "Smile-to-Bert: a pre-trained Transformer model for molecular property prediction using SMILES representations"

### 1 Methods

#### 1.1 General scheme

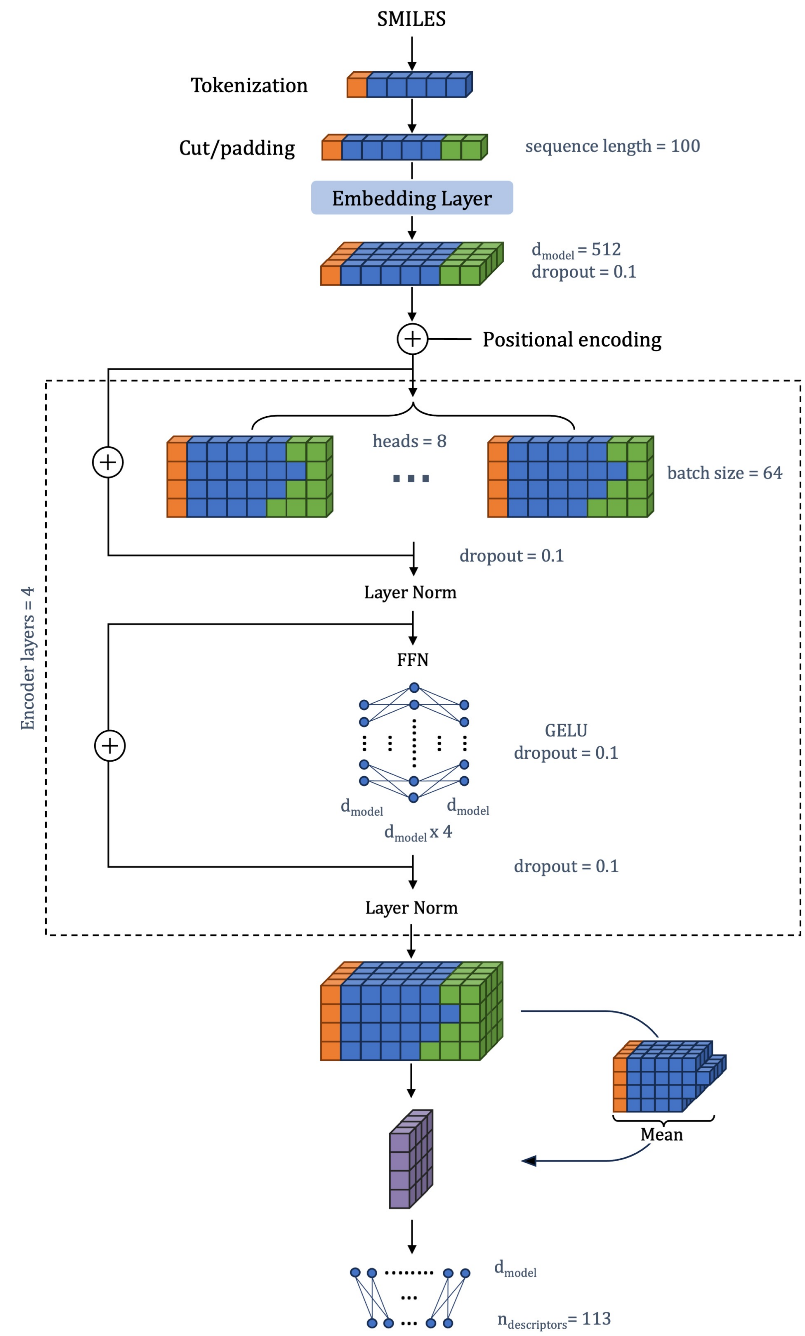


Figure S1: Scheme of the BERT architecture used to predict seven physicochemical properties using SMILES. The different parameters used are specified.

#### 1.2 Vanishing gradients problem

When training the model with scaled properties without any modification, we observed that the model was not learning. An example of this using a previous version of Smile-to-Bert with 10,000,000 samples can be seen in Figure S2. We can observe that the learning curves do not behave as expected and the values of the losses are much higher compared to the learning curves when multiplying the scaled properties by a factor of 100 (Figure S3).

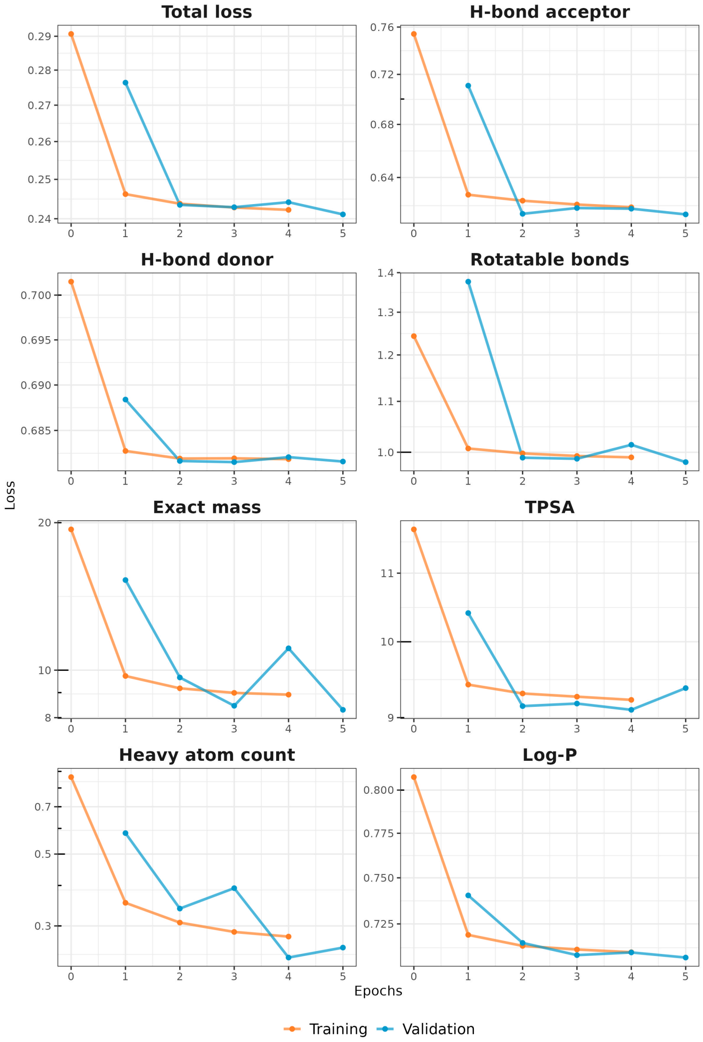


Figure S2: Learning curves when not multiplying the physicochemical properties by a factor.

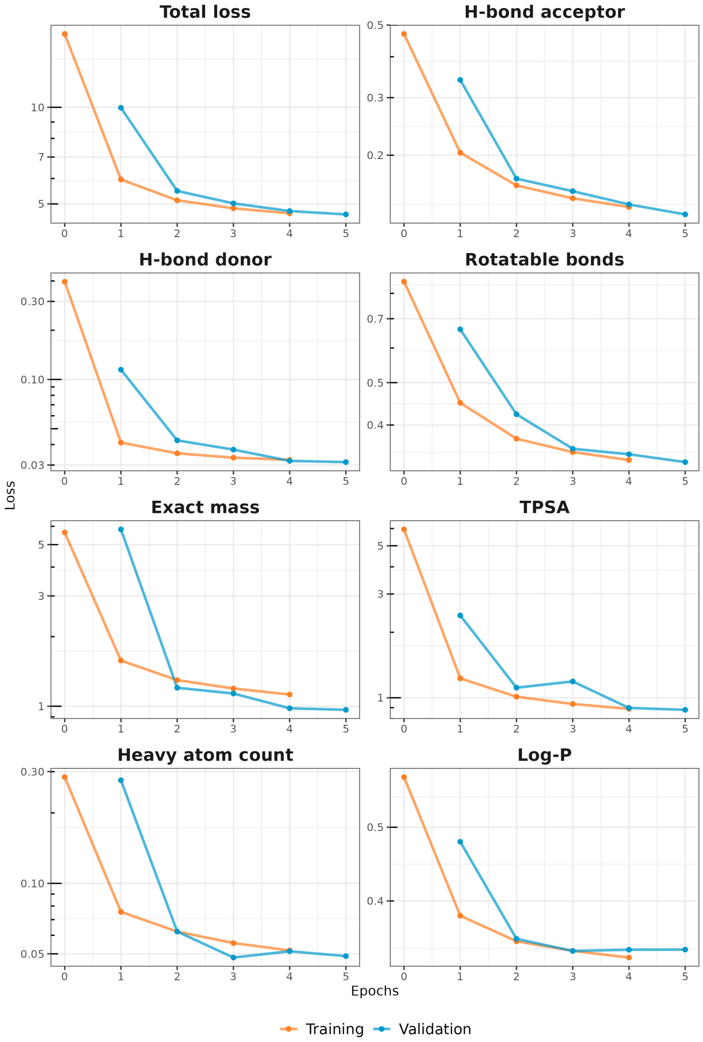


Figure S3: Learning curves when multiplying the physicochemical properties by a factor of 100.

Using Tensorboard, we obtained the distribution of the gradients for the weights of the feed forward neural network responsible of predicting the physicochemical properties for both cases (Figure S4).

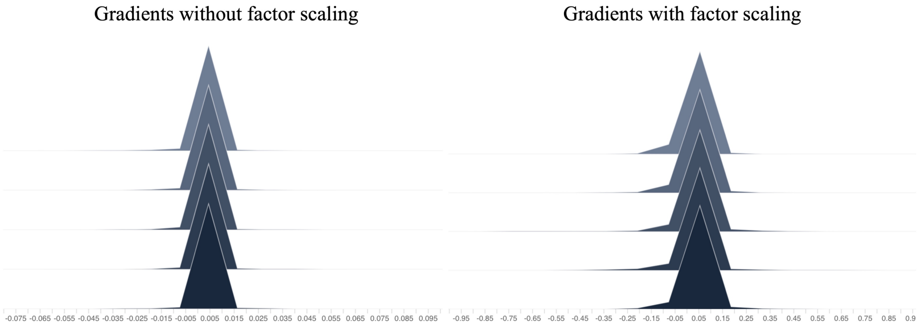


Figure S4: Gradients when not scaling and when scaling by a factor of 100.

In Figure S4, we can see that the gradients are highly distributed around zero when not scaling by a factor of 100, indicating a potential vanishing gradients problem.

#### 2 Results - Molecular descriptors learning curves

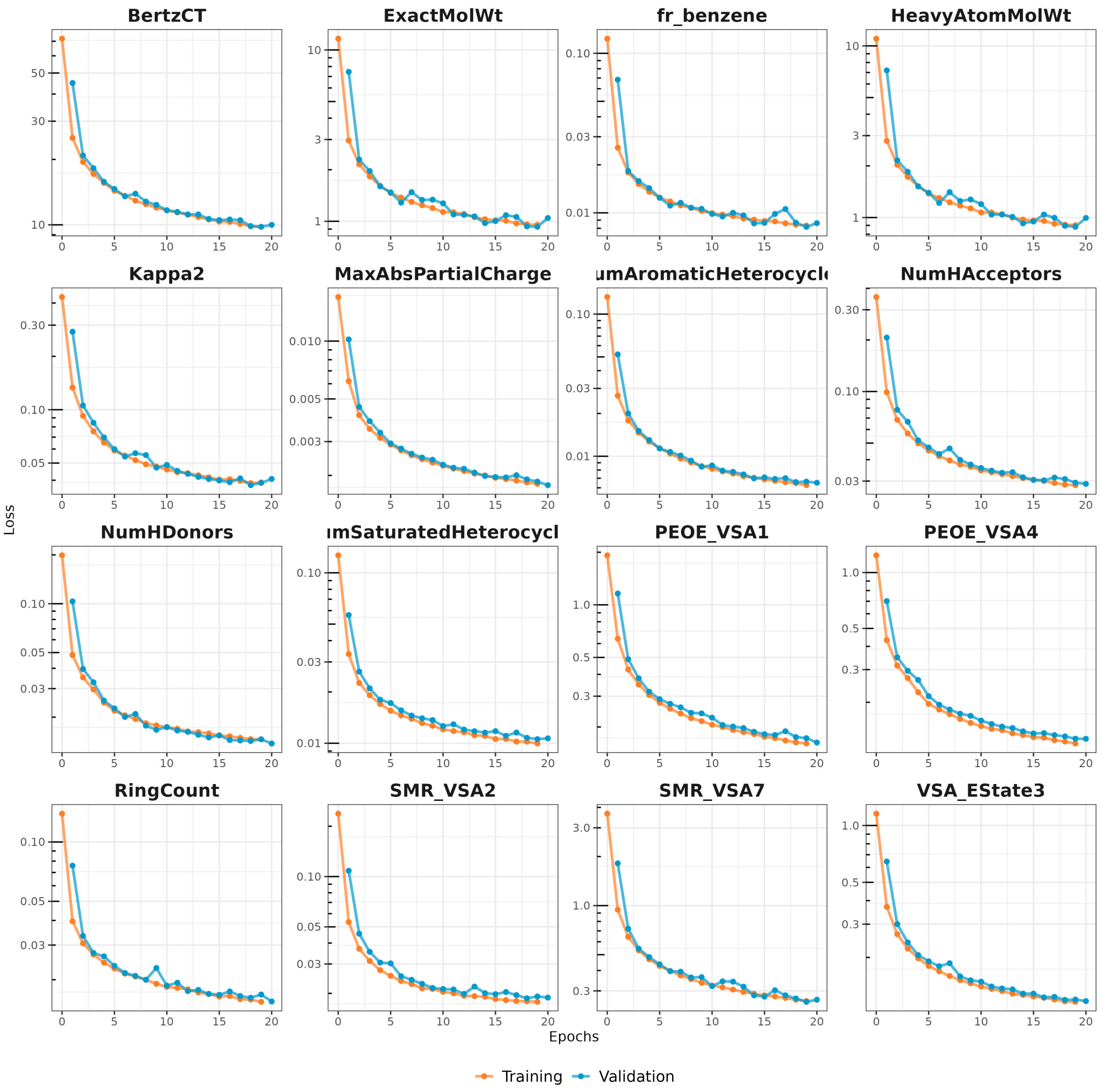


Figure S5: Learning curves for a set of molecular descriptors

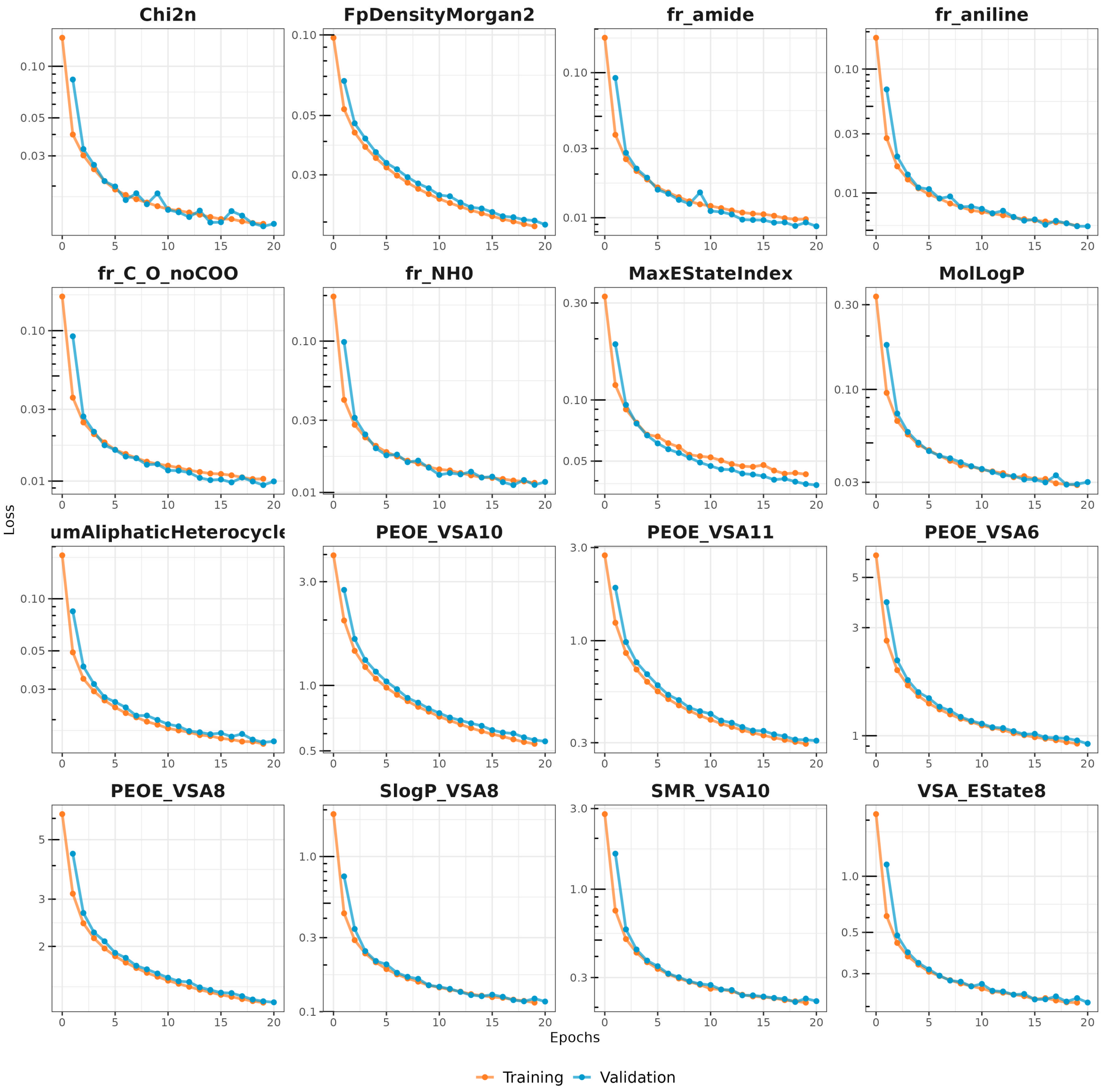


Figure S6: Learning curves for a set of molecular descriptors

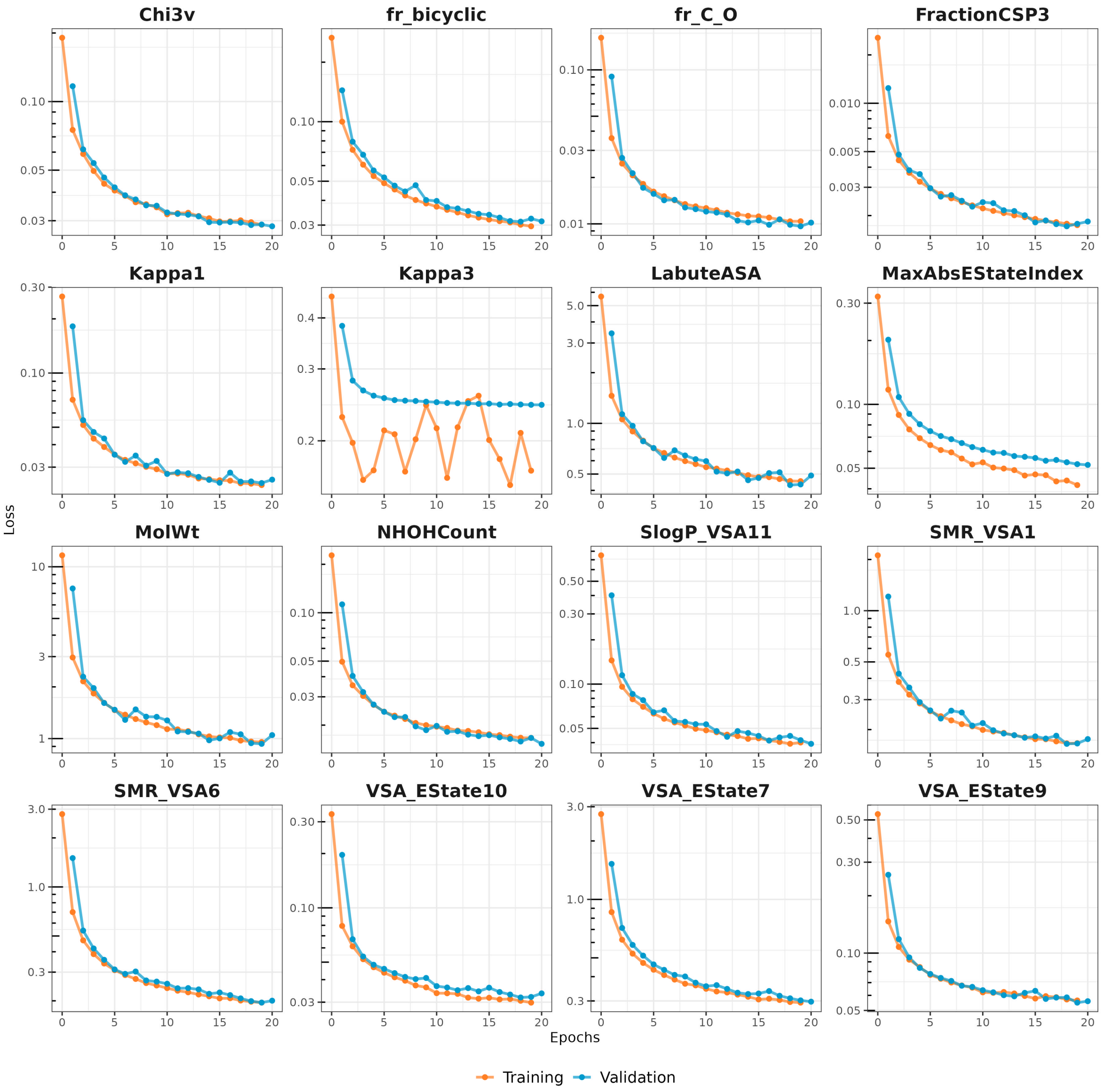


Figure S7: Learning curves for a set of molecular descriptors

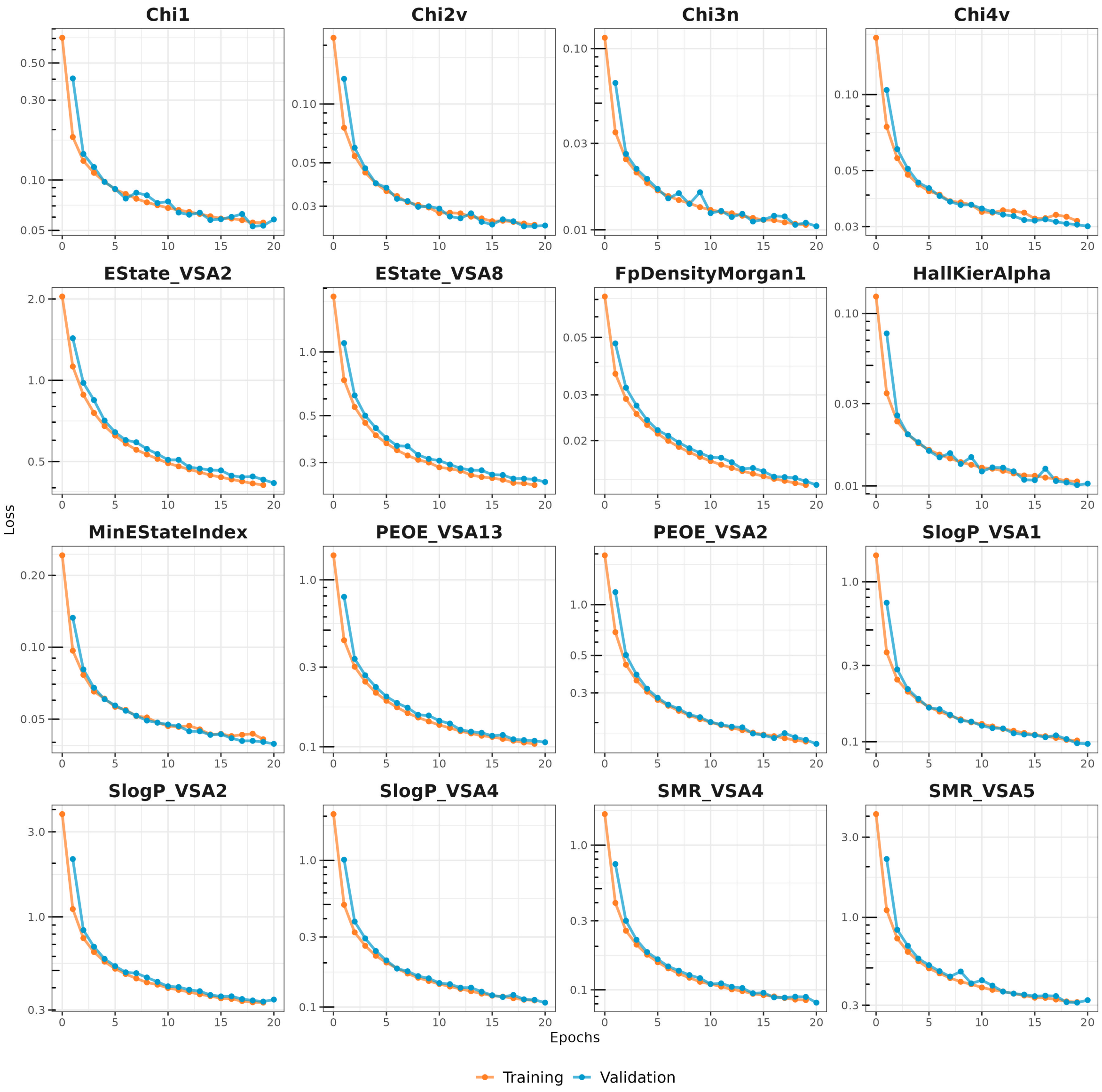


Figure S8: Learning curves for a set of molecular descriptors

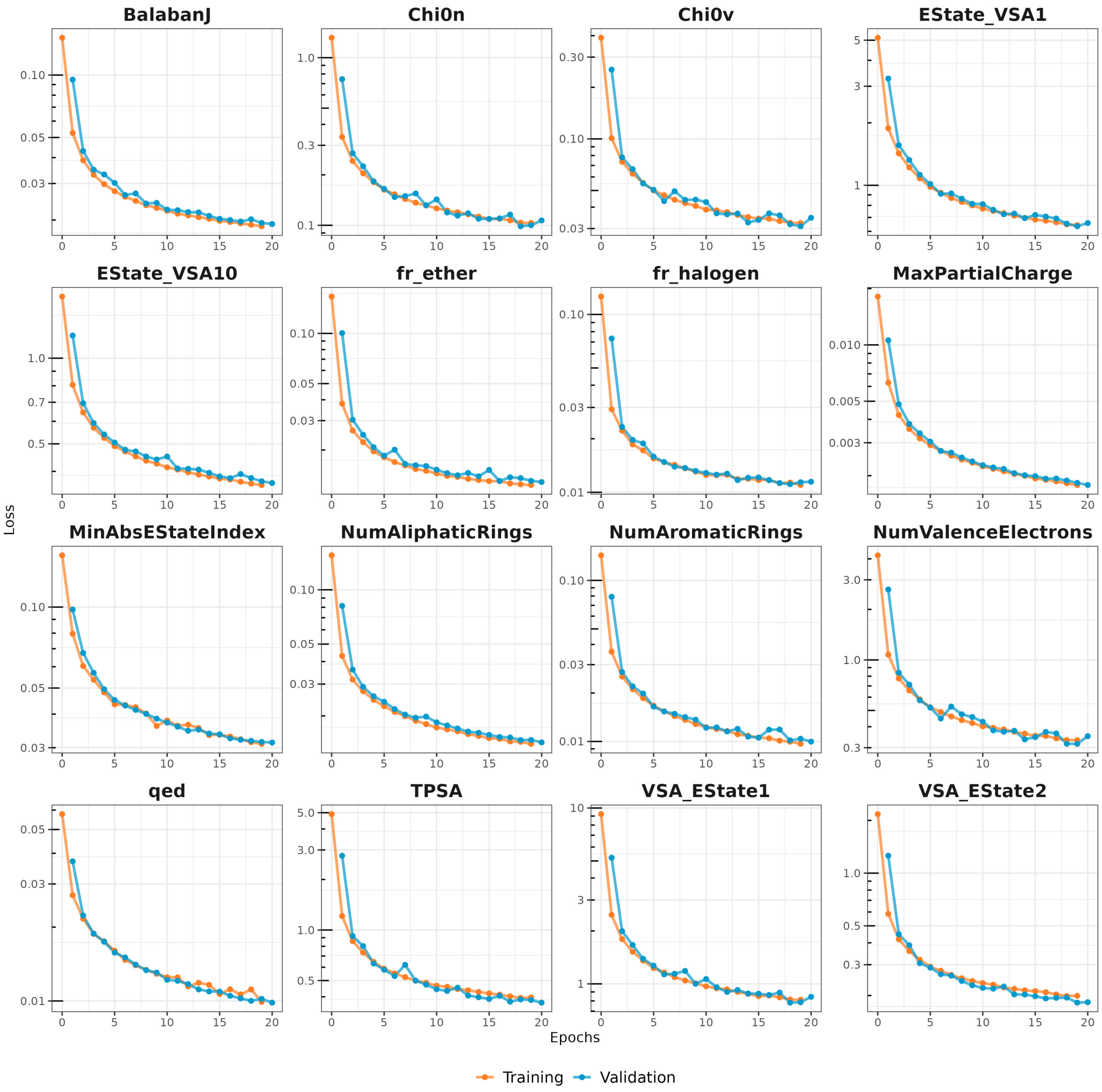


Figure S9: Learning curves for a set of molecular descriptors

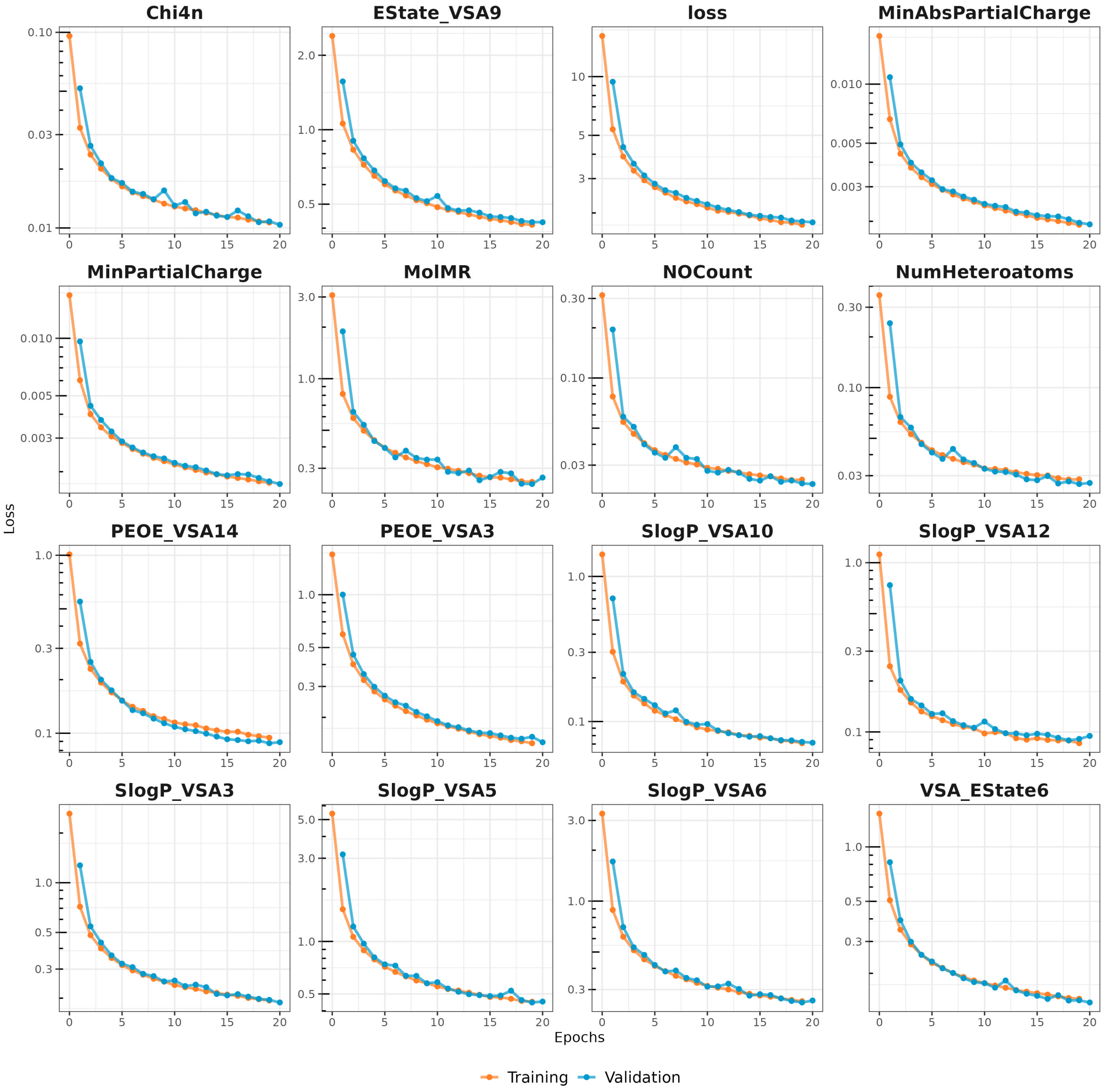


Figure S10: Learning curves for a set of molecular descriptors

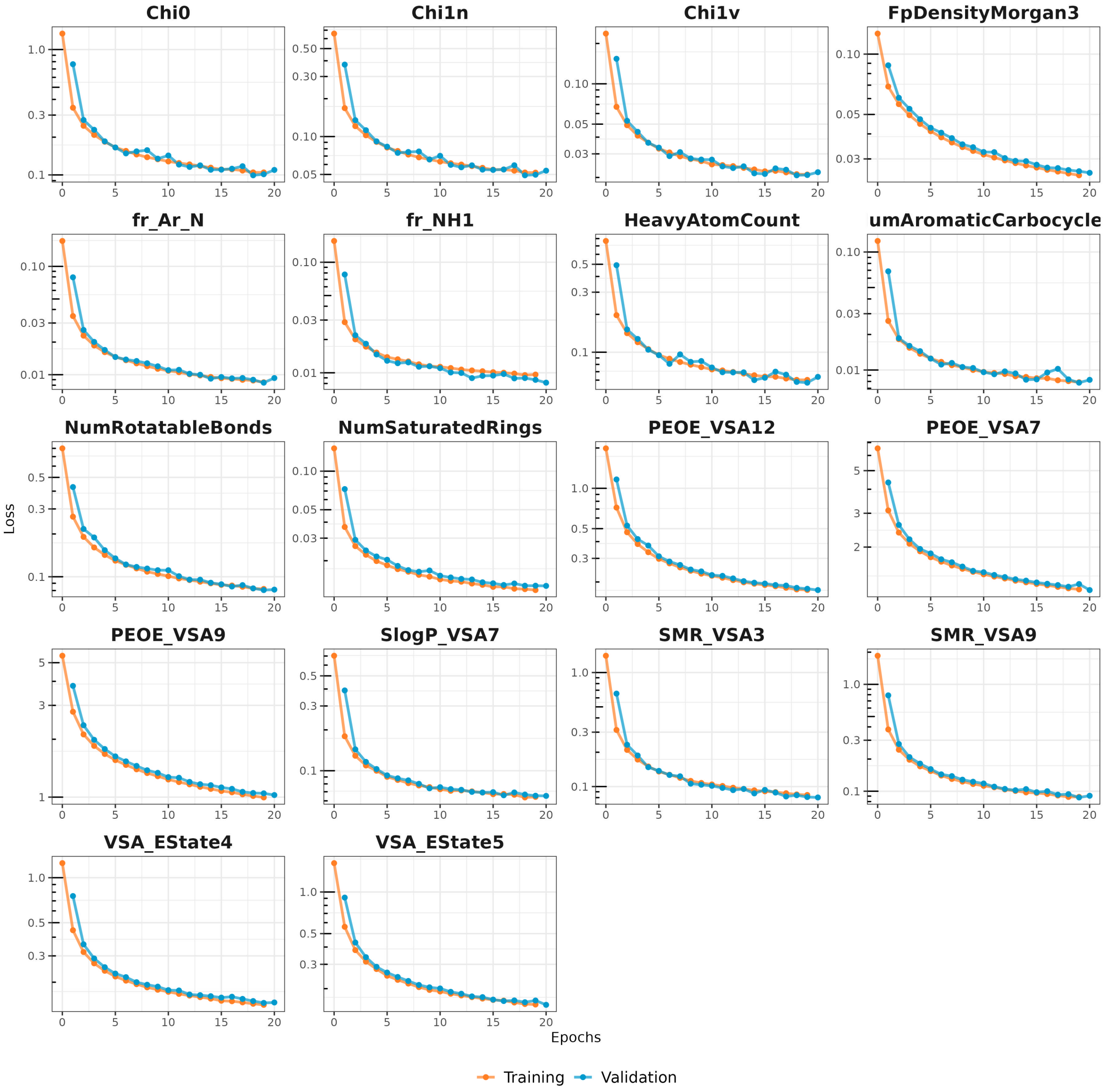


Figure S11: Learning curves for a set of molecular descriptors
